# Human Strategy Adaptation in Reinforcement Learning Resembles Policy Gradient Ascent

**DOI:** 10.1101/2025.07.28.667308

**Authors:** Hua-Dong Xiong, Li Ji-An, Robert C. Wilson, Marcelo G. Mattar

**Affiliations:** School of Psychology, Georgia Institute of Technology; Neurosciences Graduate Program, University of California San Diego; Department of Psychology, New York University

**Author notes:** Co-first authors (Hua-Dong Xiong, Li Ji-An). Co-senior authors (Robert C. Wilson, Marcelo G. Mattar).

**Keywords:** Reinforcement Learning, Decision Making, Meta-Learning, Cognitive Modeling

## Abstract

A hallmark of intelligence is the ability to adapt behavior to changing environments, which requires adapting one’s own learning strategies. This phenomenon is known as learning to learn or meta-learning. Although well established in humans and animals, a computational framework that characterizes how biological agents adapt their learning strategies through experience remains elusive. Here we posit that humans update their learning strategies online through a gradient-based meta-learning process, effectively optimizing how they learn. However, estimating how these strategies evolve over time remains a significant challenge since traditional cognitive models, such as reinforcement learning (RL), typically assume that agents use static strategies. To address this, we introduce *DynamicRL*, a method that leverages neural networks to estimate the evolution of an individual’s RL strategy by tracking cognitive parameters such as learning rates over time. Across four human bandit tasks, DynamicRL consistently outperforms traditional RL models with fixed parameters in fitting behavior, confirming that humans adapt their RL strategies over time. RL parameters estimated by DynamicRL reveal trajectories that systematically increase the expected reward of the RL strategy. The parameter updates at each step resemble policy gradient ascent, and their optimality correlates with the strength of the gradient signal. Moreover, these RL parameters evolve more slowly than decision variables, supporting the hierarchical relationship between strategy learning and value learning. Our work provides a computational framework that expands the hypothesis space from understanding strategies to understanding strategy adaptation, bridging adaptive behavior in biological and artificial intelligence through meta-learning.

## 1 Introduction

Intelligence is flexible and dynamic, shaped by evolutionary pressures in an ever-changing environment, demanding adaptation extend beyond immediate behavior to the learning process itself. This view resonates with Anderson’s general principle of rationality, which states that “the cognitive system operates at all times to optimize the adaptation of the behavior of the organism” (Anderson, 2013). In line with this principle, Harlow’s pioneering work (Harlow, 1949) and numerous subsequent studies show that humans and animals dynamically adjust their learning strategies to meet task demands, i.e., they *learn to learn* (Behrens et al., 2007; Browning et al., 2015; Nassar et al., 2010, 2012; Piray and Daw, 2020; Soltani et al., 2006; Hattori et al., 2023). For example, individuals learn to rely more on recent experience by adopting higher learning rates in rapidly changing environments, whereas they use lower learning rates in stable environments (Piray and Daw, 2021).

These adaptive principles also apply to artificial intelligence (AI), offering clues into the underlying computational mechanisms. The idea of *learning to learn* features prominently under “meta-learning”, where learning algorithms adapt flexibly to diverse tasks (Andrychowicz et al., 2016; Finn et al., 2017; Li et al., 2018; Ortega et al., 2019; Wang et al., 2017). In the case of large language models (LLMs), this similar process — known as “in-context learning” (ICL) — enables models to adapt rapidly to new examples online without model retraining (Lampinen et al., 2024), akin to human-like rapid adaptation. Recent work has shown that this flexible learning resembles a form of online, gradient-based meta-optimization of the ongoing algorithm in response to task feedback (Ahn et al., 2023; Cheng et al., 2024; von Oswald et al., 2023a,b; Coda-Forno et al., 2023).

Insights from LLMs provide a compelling clue about the principle of adaptation: the refinement of learning strategies observed in biological agents may likewise be interpreted as online, gradient-based optimization. We term this the `gradient-based meta-learning hypothesis:` that individuals engage in a meta-learning process, dynamically adapting their strategy online to improve task performance. We therefore test a series of progressive predictions from this hypothesis. First, a model capturing these dynamic strategies will better explain behavior. Second, this adaptation leads to improved performance. Finally, the adaptation process resembles gradient-based learning.

Reinforcement learning (RL) (Rescorla and Wagner, 1972; Sutton and Barto, 1998) provides a natural computational framework to test this hypothesis, as it offers a simple and well-established formalism for modeling learning from reward feedback across species and tasks. However, capturing how the RL strategies evolve over time remains a key methodological challenge. Traditional cognitive models use a small set of fixed parameters and therefore cannot capture strategy adaptation over time. Recurrent neural networks (RNNs) trained via meta-learning can only qualitatively describe biological behavior (Wang et al., 2018). Furthermore, while hierarchical Bayesian models (Mathys et al., 2011, 2014; Piray and Daw, 2021) can infer such strategy adaptation from behavior, they impose strong assumptions about the underlying generative process that may not hold for biological behaviors that deviate from optimality. This highlights a fundamental limitation in our ability to characterize the quantitative principles of biological meta-learning. The field thus lacks a robust, data-driven framework for tracking how learning strategies evolve with experience over time.

In this paper, we first address this methodological gap by introducing DynamicRL, a novel computational framework that uses neural networks to robustly estimate RL parameters on a trial-by-trial or block-by-block basis. Second, we apply this framework to test three progressive predictions of the `gradient-based meta-learning` hypothesis in human strategy adaptation across four diverse value-based decision-making tasks. We show that the DynamicRL method consistently outperforms traditional RL models with fixed parameters in fitting human behavior, indicating that participants dynamically adjusted their learning strategies throughout the task. Furthermore, these dynamically estimated RL parameter trajectories exhibit a systematic progression toward higher expected rewards, suggesting that this adaptation reflects an online (i.e., in real time) refinement of learning strategies. Critically, the direction of parameter updates broadly aligns with the policy gradient, the changes in RL parameters that most increase the expected reward, reflecting gradient-based optimization of the strategy. Moreover, the signal-to-noise ratio of the gradient correlates with the efficiency of meta-learning. Finally, we find that the meta-level adaptation of RL parameters evolves on a slower timescale than the RL process that updates decision variables, indicating a hierarchical organization of learning. Our results support the prediction of `gradient-based meta-learning` hypothesis, where individuals not only learn action values but also gradually refine their learning strategies in a manner resembling policy gradient ascent.

Taken together, our computational framework characterizes how learning strategies evolve over time, providing a principled account of strategy adaptation as gradient-based meta-learning. It expands the hypothesis space from understanding strategies to understanding strategy adaptation, and provides evidence that human strategy adaptation in RL reflects this process. This perspective offers a unified computational account of how learning strategies evolve through experience, bridging adaptive mechanisms across biological and artificial systems.

## 2 Results

### 2.1 A meta-learning perspective on the online adaptation of RL parameters

To study strategy adaptation in RL, we focus on multi-armed bandit tasks, a common paradigm in psychology and neuroscience where participants choose between several options with unknown reward distributions. The expected rewards of options may drift over time, reflecting environmental volatility. Additionally, the reward delivered on each trial is often stochastic, reflecting environmental uncertainty. Below, we formalize gradient-based meta-learning hypothesis within this context by elucidating two distinct levels of learning.

#### First-level: learning action values

To maximize rewards in the bandit task, an agent must learn the value of each action from noisy experience to make decisions (Fig. 1a). The first-level learning addresses how the agent learns the action values. This process is typically modeled using the standard Rescorla-Wagner rule, where the value of a chosen action *i* at trial *t, Q*_*i,t*_, is updated by the reward prediction error:

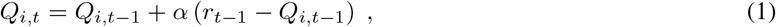

where *r*_*t*™1_ is the reward received on the previous trial and *α* governs how much weight is given to the most recent outcome.

**Figure 1.**
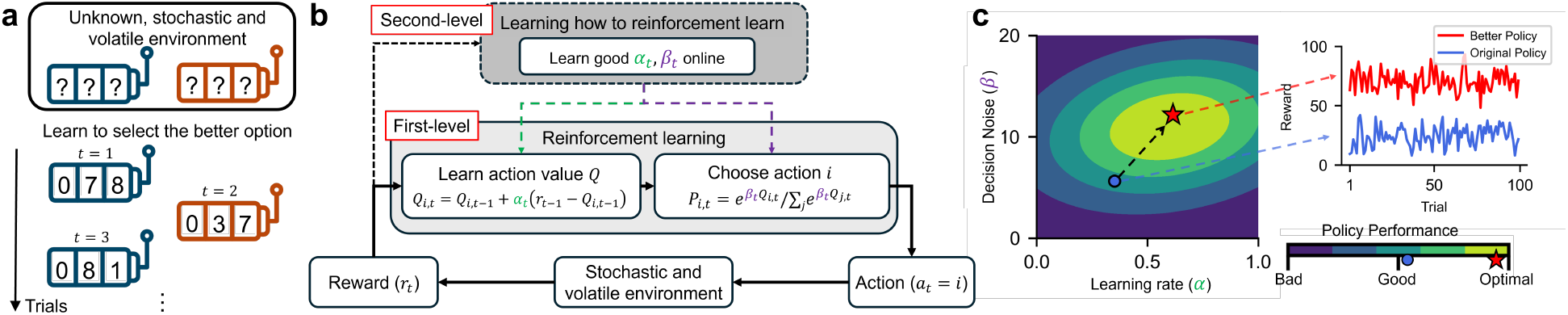
Meta-learning in a two-armed bandit task. **a**, Illustration of a two-armed bandit task. Participants interact with two slot machines with unknown expected rewards and learn to estimate the value of each machine through repeated choices. **b, Meta-learning schematic:** at the first level, agents take actions, interact with the environment, receive rewards, and update internal states according to an RL model; at the second level, agents learn to adjust their RL parameters (*α*_*t*_, *β*_*t*_) online to improve task performance. **c**, A conceptual “reward landscape” over RL parameters illustrates how different settings of *α* and *β* yield different expected rewards. The agent begins with a policy (strategy) using suboptimal parameters (blue dot), evaluates reward feedback, and updates RL parameters in a direction that improves task performance. Over time, it progresses toward a better policy (red star). **Upper-right inset:** (hypothetical) rewards obtained by the original and the better strategies, demonstrating the advantage of using better RL parameters.

To translate these learned values into actions, a canonical approach uses the softmax function, which balances the exploitation of the highest-value option with the exploration of alternatives:

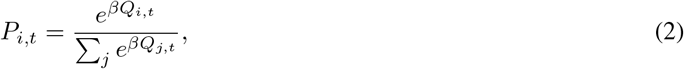

where the inverse temperature *β* controls the degree of decision stochasticity. Together, *α* and *β* determine the agent’s RL strategy. We focus on this two-parameter RL model for its simplicity and normality.

#### Second-level: learning to adapt RL parameters

The choice of these RL parameters is critical for task performance. For a given task, any combination of *α* and *β* defines an RL strategy that generates a specific expected reward. The set of all combinations thus form the “reward landscape” (Fig. 1c). Existing cognitive models typically adopt fixed parameters (i.e., a single position on the reward landscape), implicitly assuming that the agent uses a constant strategy throughout the experiment.

We posit that rational agents (such as humans) engage in a second-level meta-learning process that learning not only learning action values but also learning how to learn by adapting their RL parameters. Since the optimality of RL parameters depends on environmental factors such as volatility and stochasticity (Piray and Daw, 2021), which are unknown to the agent a priori, this adaptation must occur online. This second-level learning can be viewed as an online optimization within the parameter space of *α* and *β*. Rewards provide feedback signals that enables the agent to estimate the local gradient on the reward landscape and adjust its RL parameters accordingly. This forms our gradient-based meta-learning hypothesis: through experience, humans gradually refine their RL parameters in a direction that aligns with gradients to improve task performance, in a manner resembling online gradient-based meta-learning.

### 2.2 DynamicRL better explains human behavior by modeling strategy adaptation

To test our hypothesis, we analyze human behavior across four diverse multi-armed bandit tasks. These tasks cover a broad range of environmental designs, including differences in the number of options, reward volatility, stochasticity, and block length (see Sec. 4.1 for details). They fall into two widely studied categories: stationary bandit tasks featuring stable reward distributions with no environmental volatility, and restless bandit tasks involving dynamically shifting rewards that demand continual adaptation. For the restless bandit tasks, we analyze `Bahrami2020` (Bahrami and Navajas, 2020) and `Suthaharan2021` (Suthaharan et al., 2021) datasets. `Bahrami2020` involves a 4-armed bandit task in which each option’s expected reward drifts over time (Fig. 2b). `Suthaharan2021` features a 3-armed probabilistic reversal learning task, where choice probabilities change abruptly and unpredictably. For the stationary bandit tasks, we analyze `Gershman2018` (Gershman, 2018) and `Wilson2014` (Wilson et al., 2014) datasets. `Gershman2018` includes a 2-armed bandit task with Gaussian rewards with mean resampled in each block. `Wilson2014` presents a 2-armed bandit task with varying block lengths and differing numbers of forced-choice trials to manipulate uncertainty.

**Figure 2.**
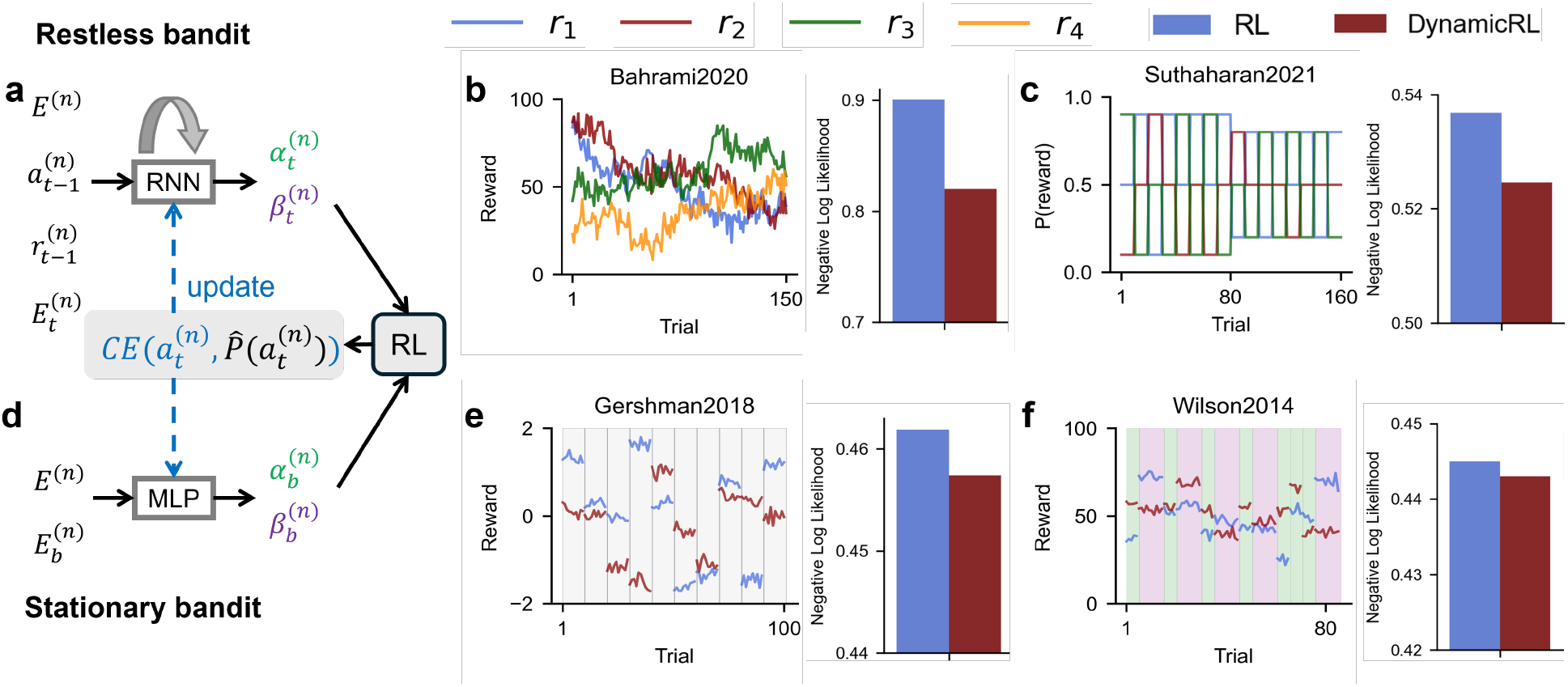
DynamicRL better explains behavioral variability across four tasks. **a, d**, Schematic of the `DynamicRL`. The model estimates time-varying RL parameters, which are then fed into an RL model to predict the human’s next choice. The resulting cross-entropy (CE) loss updates the neural network. **a**, For restless bandit tasks, an RNN estimates RL parameters at the trial level. Inputs consist of a participant-specific embedding *E*^(*n*)^, the previous action 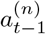 and reward 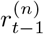, and trial-level task information 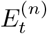. **d**, For stationary bandit tasks, a two-layer multilayer perceptron (MLP) estimates RL parameters at the block level using the participant embedding *E*^(*n*)^ and trial-level task information 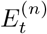 as input. **b, c, e, f**, Multi-armed bandit tasks and model fitting performance. **Left panels**: Reward structures for the four datasets. The expected reward of each option varies across trials within a block for the restless bandit tasks (**b, c**), and it is fixed within a block for the stationary bandit tasks (**e, f**). **b**, `Bahrami2020`: a 4-armed restless bandit task where participants choose among actions *A*_1_–*A*_4_, each with rewards ranging from 0 to 100 that evolve over time via an Ornstein–Uhlenbeck process centered at 50. **c**, `Suthaharan2021`: a 3-armed probabilistic reversal learning task where participants choose among actions *A*_1_–*A*_3_, each associated with binary rewards and time-varying reward probabilities. **e**, `Gershman2018`: a 2-armed bandit task where the expected rewards for the two options are sampled from a hierarchical Gaussian distribution. **f**, `Wilson2014`: a 2-armed bandit task where the difference in expected rewards between arms is sampled from a predefined set. The first 4 choices in each block are forced, with block horizons of either 1 (green shading) or 6 (purple shading). For visualization purposes, the reward standard deviation for `Gershman2018` and `Wilson2014` has been divided by four. **Right panels:** On unseen test data, the `DynamicRL` model outperforms classic RL models with fixed parameters (lower negative log-likelihood indicate better fit).

A key challenge in testing whether humans dynamically adjust their RL strategies is the difficulty of estimating cognitive parameters over time from limited data, as each trial often provides only a single data point per participant. To address this, we develop the `DynamicRL` method (see Sec. 4.2 for technical details), which leverages a neural network to estimate dynamic RL parameters over the course of an experiment.

The neural network architecture in DynamicRL is designed to flexibly accommodate the temporal structure of tasks. In restless bandit tasks (`Bahrami2020`, `Suthaharan2021`), the expected reward of each option fluctuates across trials (*t*) due to volatility. We therefore define the RL strategy as dynamic across trials, parameterized by trial-by-trial RL parameters *α*_*t*_ and *β*_*t*_. To estimate these trial-level parameters for each participant *n*, we use a recurrent neural network (RNN) (with gated recurrent units (Cho et al., 2014)). The inputs to this RNN include the previous action 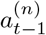, previous reward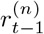, participant information *E*^(*n*)^, and trial-level task information 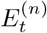(Fig. 2a, see Sec. 4.2.1 for details).

By contrast, in the stationary bandit tasks (`Gershman2018`, `Wilson2014`), the expected value of each option remains constant across trials within a block, yet differs across blocks (Fig. 2e,f). Accordingly, we define RL strategies with parameters 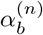 and 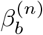 varying at the block level. These parameters are estimated solely from block-level task information 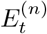 via a two-layer perceptron (Fig. 2d, see Sec. 4.2.1 for details). This formulation allows us to examine RL strategy adaptation on a slower, block-level timescale. Consequently, trial-by-trial actions and rewards are not used in the estimation process.

We next evaluated whether `DynamicRL` can better explain human behavior than competing models through a formal model comparison using nested cross-validation (see Sec. 4.2.2 for details). `DynamicRL` consistently outperformed RL models with fixed parameters across all four datasets, achieving a lower test negative log-likelihood, indicating superior predictive accuracy (Fig. 2b,c,e,f). We then compared `DynamicRL` to an alternative approach for estimating dynamic RL parameters: fitting static RL models on adjacent trials within a sliding window. However, such methods typically perform poorly and are prone to overfitting due to the limited data available in each window (see details in Sec. A.2). `DynamicRL` leverages modern deep learning training pipelines (Xiong et al., 2025a) and pools data across all participants to generate robust and reliable parameter estimates. Finally, to validate DynamicRL using data with known ground truth, we simulated RL agents with predefined parameter trajectories (constant, linearly increasing, or linearly decreasing for both *α* and *β*). `DynamicRL` accurately recovered the ground-truth parameters from the synthetic data (see Sec. A.1). The better predictive performance of `DynamicRL` demonstrates that incorporating temporal dynamics in RL parameters yields a more accurate account of human behavior. These results support the view that humans do not rely on a fixed strategy but continuously adapt their learning parameters throughout the experiment, confirming our first prediction of `gradient-based meta-learning` hypothesis.

### 2.3. Humans improve their strategies by adjusting RL parameters online

We next examined the trajectories of these RL parameter changes over time. The `DynamicRL` method revealed systematic temporal patterns in both learning rates *α* and inverse temperatures *β* (averaged over participants; see Fig. 3). In the restless bandit tasks, `Bahrami2020` (Fig. 3a) and `Suthaharan2021` (Fig. 3b), both parameters gradually increase over trials. This shows that humans increasingly filter past information in volatile environments. Notably, in blocks 2–4 of `Suthaharan2021`, RL parameters plateau more quickly, suggesting faster adaptation as participants become familiar with the task structure. In the stationary bandit tasks, `Gershman2018` and `Wilson2014`, we also observe systematic changes in *α* and *β* across blocks (Fig. 3c,d). These findings again corroborating that humans continuously adjust their RL strategies at both trial and block levels.

**Figure 3.**
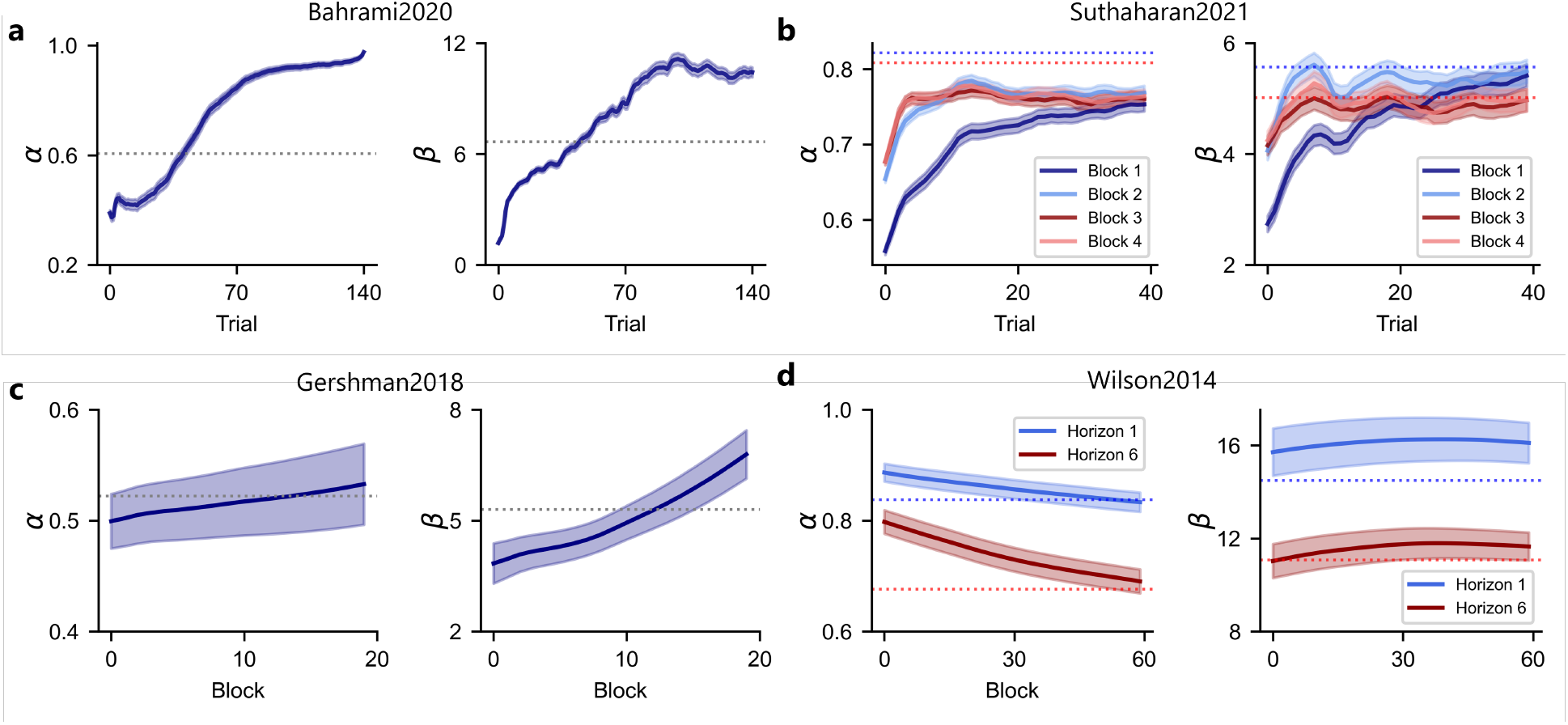
DynamicRL reveals systematic adaptations in human RL parameters across four tasks. **a–d**, Solid lines show RL parameters (*α* and *β*) estimated by `DynamicR`L, averaged across participants over the course of each experiment. Shaded areas indicate 95% confidence intervals. Dotted lines represent averaged RL parameters estimated from traditional RL models with fixed parameters.

How can we interpret the observed changes in RL parameters? The second prediction of the *gradient-based meta-learning* hypothesis is that participants adapt their RL parameters to improve performance. Thus, they should perform better in the later stages of the experiments. We first conducted a model-agnostic analysis comparing within-subject behavioral performance between the first and second halves of each experiment, while controlling for task conditions using a mixed-effects model (see Sec. 4.3 for details). We found significant improvements in second-half performance relative to the first half in `Bahrami2020`, *t*(950) = 18.14, *p*.001; `Suthaharan2021`, *z* = 5.37, *p*.001; `Gershman2018`, *t*(43) = 2.34, *p* =.024; and `Wilson2014`, *z* = 4.736, *p*.001. These results confirm that participants’ performance improved systematically over time.

To understand how participants improve their behavioral performance by adapting RL parameters, we overlaid the RL parameter trajectories (averaged across participants) onto the “reward landscape,” which shows the expected reward 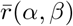for each possible parameter combination (Fig. 4). We generated reward landscape by simulating RL agents with different combinations of the learning rate (*α*) and inverse temperature (*β*) to compute the expected reward for each RL strategy (see Sec. 4.4 for details). We observed a clear and consistent pattern across all four datasets: the averaged RL parameters defining the strategy evolve from lower-reward to higher-reward regimes over the course of the experiment (white-to-red curves in Fig. 4). We further visualized the distribution (kernel density) of RL parameters across participants at the beginning and end of the experiment, revealing a shift from lower-reward to higher-reward regimes (dashed contours in Fig. 4).

**Figure 4.**
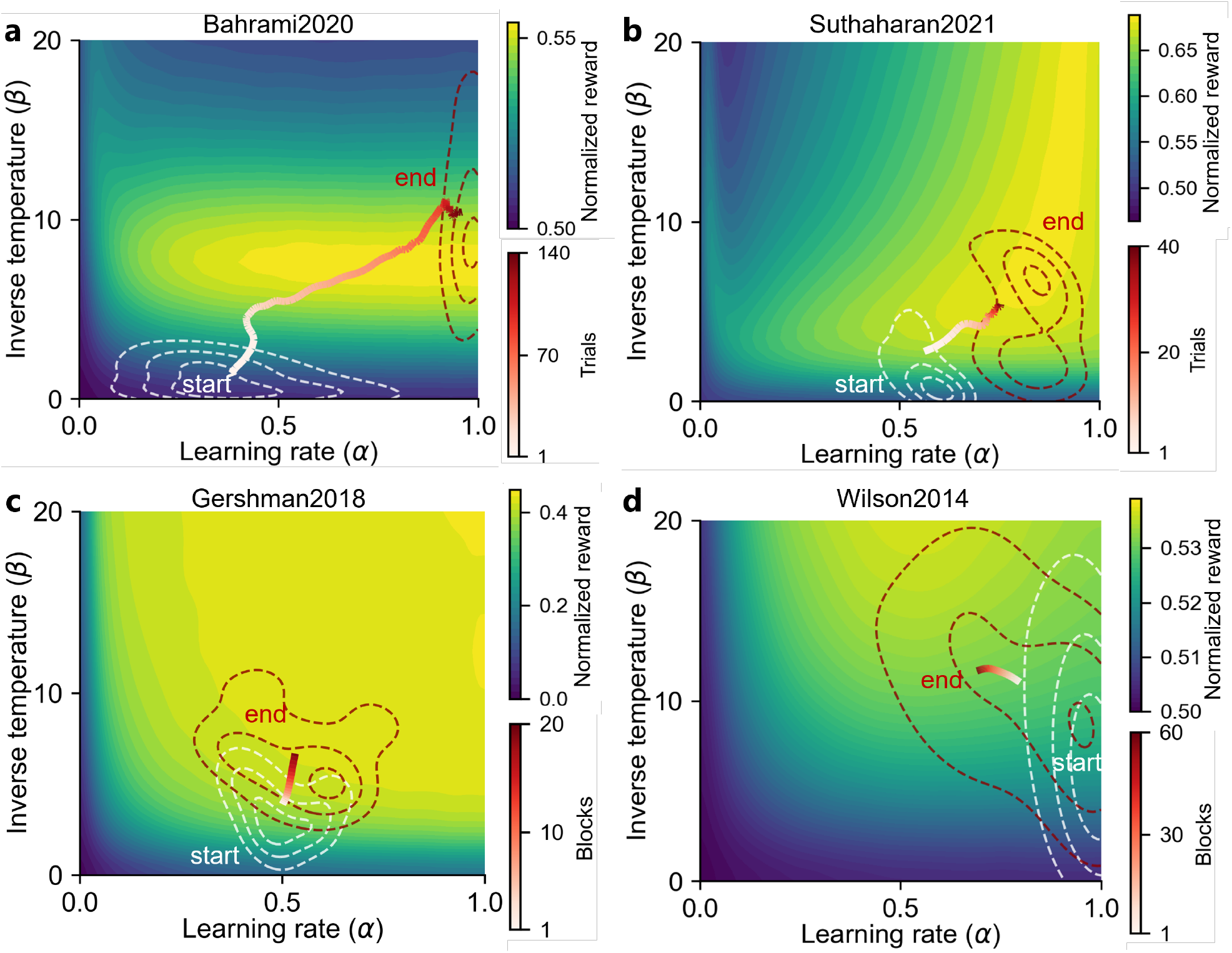
Trajectories of RL parameters over the course of the experiment. This figure illustrates the evolution of participants’ RL strategies across four tasks. Each panel displays the reward landscape for a task, defined over a two-dimensional RL parameter space parameterized by learning rate (*α*) and inverse temperature (*β*) for a task. Colored contours indicate the expected normalized reward for each parameter combination, computed from simulations of RL agents on the task. Dashed contours represent kernel density estimates of participants’ dynamic RL parameters, as estimated by `DynamicRL`, at the start (white) and end (dark red) of the experiment. The white-to-red solid lines trace the average temporal evolution of participants’ estimated RL parameters. This average trajectory toward brighter, higher-reward regions demonstrates that participants adaptively optimized their RL strategies as the experiment progressed.

These analyses suggest that participants improved their behavior over time, an improvement associated with the online adaptation of RL parameters toward more effective strategies. This supports our second prediction of the *gradient-based meta-learning* hypothesis.

### 2.4. RL parameters adaptation resembles policy gradient ascent

Having shown that humans adapt their learning strategies to improve performance, we next examine our third prediction: whether this strategy adaptation process is analogous to policy gradient ascent — a basic form of gradient-based learning algorithms in RL. Policy gradient estimates reward gradient, how changes in parameters affect expected rewards of strategy and updates parameters accordingly to maximize performance. Visual inspection of the humans’ parameter trajectories suggests they broadly follow the reward gradients (Fig. 4); here, we quantify this alignment using two complementary metrics.

First, as a measure of directional optimality, we calculate cosine similarity between the estimated RL parameter update at each step and the optimal gradient derived from the reward landscape (Fig. 5a; see Sec. 4.5.1 for details). In artificial agents, policy gradient ascent typically requires averaging over many repeated episodes to reduce the variance of gradient estimation. However, each human participant completed only a single action-outcome sequence per task. This limited data naturally results in high variance in estimated gradient signals, so the RL parameter updates are expected to be noisy as they can only depend on several adjacent trials. Consequently, we do not expect perfect alignment with the optimal gradient, we hypothesize that at the population level, RL parameter updates should exhibit a positive cosine similarity with it. As predicted, we find a positive alignment at the population level across four tasks, with the distribution of cosine similarity values skewed significantly above zero (Fig. 5b).

**Figure 5.**
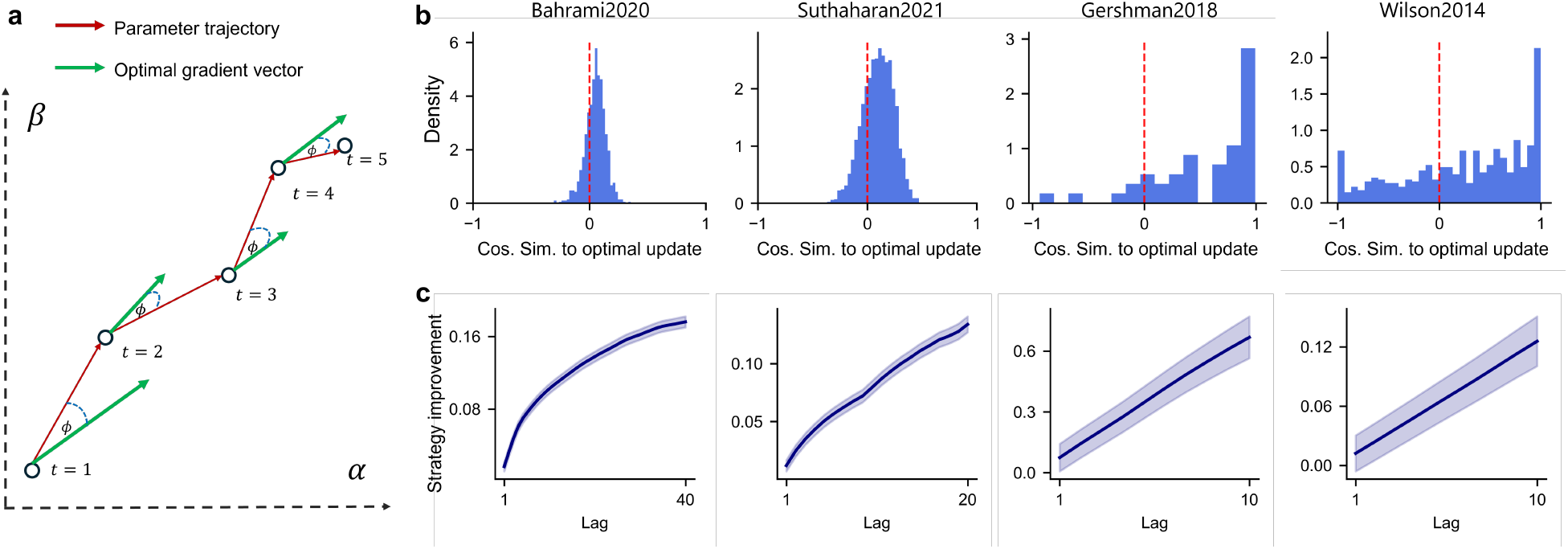
Human meta-learning updates align with optimal updates. **a**, Schematic illustrating cosine similarity between human RL parameter updates and the optimal gradient direction. Each circle represents an update step; the red arrow indicates the RL parameter update, and the green arrow shows the optimal gradient direction. *θ* indicates the similarity between the RL parameter update and the optimal gradient ascent direction. **b**, Distribution of cosine similarity for single-step parameter updates, computed from dynamically estimated RL parameters, showing a statistical alignment with the optimal gradient. **c**, Effect size of the alignment with the optimal gradient. The effect size increases with larger window sizes, indicating a consistent improvement in RL strategies.

While directional alignment is intuitive, it is sensitive to the numerical scales of the RL parameters (*α* and *β*). Therefore, the degree of optimality it reflects also depends on the parametrization of the model, as the relative scaling of the parameters is not uniquely defined. To provide a more direct and scale-independent test, we introduce a second metric, *strategy improvement*, which measures how much the expected reward of the strategy changes as participants update their RL parameters. Formally, this corresponds to the change in expected reward between the current policy and the one a few steps (“Lag”) earlier (i.e.,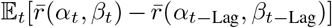). We reported its effect size (Cohen’s d), where 0 indicates no improvement and positive values indicate improvement in obtained rewards (see Fig. 5c and Sec. 4.5.2 for details). We found that strategy improvement at Lag=1 was significantly positive across all four datasets (`Bahrami2020`: *t*(143,784) = 14.15, *p* <.001; `Suthaharan2021`: aggregated across all blocks, *t*(157,871) = 4.48, *p* <.001, `Gershman2018`: *t*(835) = 10.72, *p*<.001; `Wilson2014`: Horizon 1, *t*(14,615) = 32.01, *p* <.001; Horizon 6, *t*(14,615) = 16.63, *p* <.001; *p*-values from permutation tests), showing participants’ parameter updates reliably improved their strategies. Moreover, the magnitude of strategy improvement increased with larger lag values, indicating that these improvements accumulate systematically over longer timescales.

Taken together, the positive alignment in the parameter space and the consistent strategy improvement in the reward space provide converging evidence that human strategy adaptation is functionally analogous to the policy gradient, consistent with our third prediction of *gradient-based meta-learning* hypothesis.

### 2.5 Task gradient structure informs meta-learning efficiency

In artificial learning systems, gradient-based optimization performs more efficiently when the signal-to-noise ratio (SNR) of the gradient estimates is high, as strong and reliable gradient signals enable effective learning. If human strategy adaptation relies on gradient-based mechanisms, then updates initiated at parameter positions with higher gradient SNR should produce greater strategy improvements. Therefore, we examined how the reward gradient SNR relates to strategy improvement at each step along a participant’s trajectory.

We estimated the point-wise gradient SNR on each task’s reward landscape as the ratio of the average policy gradient to its standard deviation across multiple simulations (see Section 4.6 for details). Across all four datasets, gradient SNR was significantly positively correlated with the magnitude of strategy improvement (`Bahrami2020`: *r* = 0.23, *p* <.001; `Suthaharan2021`: aggregated across all blocks, *r* = 0.54, *p* <.001; `Gershman2018`: *r* = 0.53, *p* <.001; `Wilson2014`: Horizon 1, *r* = 0.25, *p* <.001; Horizon 6, *r* = 0.23, *p* <.001; *p*-values are based on permutation tests, see Fig. A.4). We find that when the gradient signal is stronger, individual updates more reliably increase the expected reward of the strategy. This relationship further supports that humans leverage gradient information for meta-learning.

Modern gradient-based optimization algorithms, such as Adam (Kingma and Ba, 2017), adjust the step size of updates according to the reliability of the gradient estimate. Do humans also modulate their update magnitude based on gradient SNR, potentially taking larger steps in high-SNR regions? We examined the relationship between gradient SNR and the *norm* of the parameter update, representing the step size. We find that the effect of gradient SNR on update magnitude (step size) is mixed; the correlation between gradient SNR and the norm of the parameter update varies across datasets (`Bahrami2020`: *r* = 0.26, *p* <.001; `Suthaharan2021`: *r* = − 0.12, *p* <.001; `Gershman2018`: *r* = − 0.12, *p* <.001; `Wilson2014`: Horizon 1, *r* = 0.01, *p* =.457; Horizon 6, *r* = − 0.04, *p* <.001; *p*-values are based on permutation tests, see Fig. A.4). These findings suggest that while humans leverage gradient direction for strategy improvement, the regulation of update magnitude may reflect a subtler, context-dependent form of meta-control.

### 2.6 Two levels of learning show different timescales

Our gradient-based meta-learning framework assumes a two-level hierarchical learning process: a fast RL process that learns action values, and a slower meta-learning process that adapts the parameters (*α, β*) of the first level. This raises the question: can distinct learning timescales be observed in human behavior?

We conducted temporal autocorrelation analyses only on trial-level data from the restless bandit tasks (`Bahrami2020` and `Suthaharan2021`), since our *DynamicalRL* model estimates parameters on a trial-by-trial basis, allowing both first-level and second-level learning to unfold at the same temporal resolution. Our model itself is agnostic to timescales; it simply optimizes for the best fit to behavior, so any differences in learning timescale emerge directly from the data. In contrast, for stationary bandit tasks (`Gershman2018`, `Wilson2014`), RL parameters are estimated blockwise, which inherently imposes a slower timescale on first-level learning and obscures analysis of temporal dynamics. We predicted that the RL parameters (*α* and *β*) would be more stable over time (i.e., exhibit slower autocorrelation decay) than trial-by-trial variables such as environmental rewards (reflecting environmental structure) and action probabilities (logits, reflecting choice probabilities). Our results confirm this temporal hierarchy (Fig. 6). In both datasets, the learning rate (*α*) is the most stable parameter, evolving more slowly than the inverse temperature (*β*), which itself evolve more slowly than the choice-related logits.

**Figure 6.**
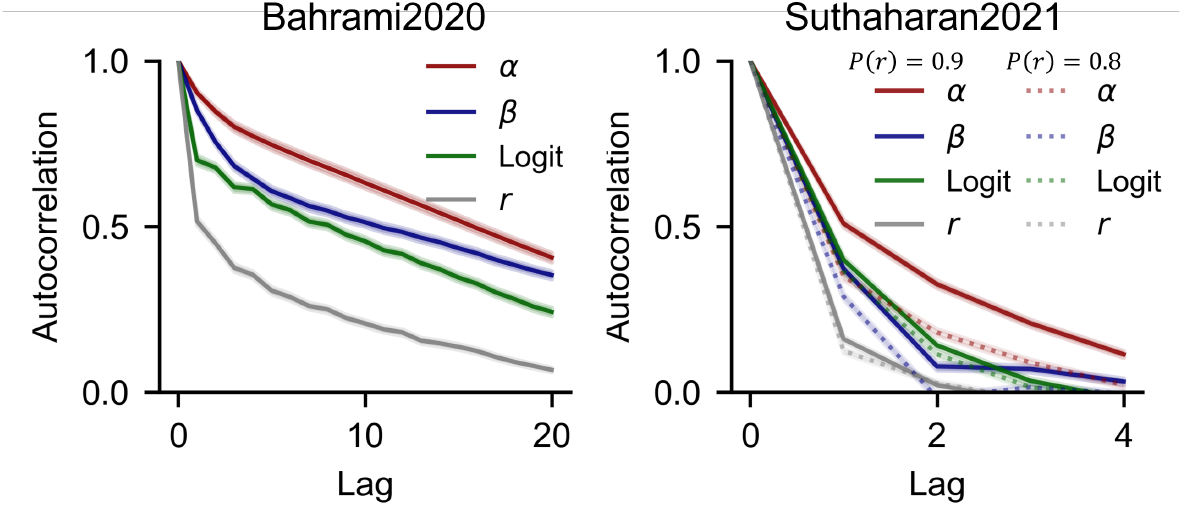
Timescale of human meta-learning processes. Autocorrelation analysis of learning rate *α*, inverse temperature *β*, logits and reward *r* reveals distinct temporal dynamics. For `Suthaharan2021`, solid and dotted lines denote two experimental conditions based on the maximum reward probability (*P* (*r*)) of the options.

We next examine whether these learning timescales reflect rational adaptation in human behavior. A rational agent should calibrate the speed of its strategy adaptations to task dynamics. The timescales of *α* and *β* reflect an agent’s belief about how often its internal model of environment should be updated. Updating too quickly leads to chasing noise, whereas updating too slowly causes the agent to miss meaningful environmental changes. We therefore tested whether individual differences in parameter timescales correlate with overall task performance (see Sec. 4.7 for details). In `Bahrami2020`, the timescale of *α* did not significantly correlate with total reward (*r* = 0.037, *p* = 0.256), whereas *β* showed a significant positive correlation (*r* = 0.256, *p* <.001). This suggests that flexible adaptation of inverse temperature (or exploration) to task dynamics is important for performance. In `Suthaharan2021`, we observed modest yet significant negative correlations between total reward and the timescales of both parameters: *α* (*r* = − 0.063, *p* = 0.004) and *β* (*r* =− 0.312, *p* <.001). These results indicate that participants with more adaptive (i.e., faster-changing) RL parameters tend to perform better, particularly when *β* adjusts more rapidly. Finally, in `Suthaharan2021`, we compared RL parameter timescales across conditions with differing reward probabilities. We found that participants updated their strategies significantly faster in the more informative, high-reward condition than in the lower-reward condition. This effect was significant for both *α* (*t*(2023) = 27.965, *p* <.001, Cohen’s *d* = 0.617) and *β* (*t*(2023) = 16.049, *p* <.001, Cohen’s *d* = 0.384). This finding further supports the idea that humans flexibly modulate the timescale of meta-learning, adjusting not only their strategies but also the speed of those adjustments in response to task structure.

## 3 Discussion

Intelligence exhibits flexibility in adapting to diverse task demands, yet how such flexibility arises — how learners adjust their learning strategies — remains poorly understood. In this work, we address this fundamental gap in two parts. Methodologically, we introduced `DynamicRL`, a novel computational approach for characterizing how learning strategies, defined by cognitive parameters, evolve with experience. Scientifically, building on this method, we proposed the *gradient-based meta-learning* hypothesis, which posits that humans adapt their learning strategies online through a process resembling policy gradient ascent. Across four diverse decision-making tasks, we found three progressive lines of evidence supporting this hypothesis. First, we demonstrated that, by modeling strategy adaptation, DynamicRL provides a better account of human behavior than traditional models with fixed parameters. Second, we observed that the trajectories of RL parameters systematically evolved toward more optimal, high-reward regions of the strategy space, leading to improved performance. Third, and most critically, we found that the direction of these strategy updates was significantly aligned with the reward gradient; a higher signal-to-noise ratio in the gradient corresponded to more efficient parameter updates, providing strong evidence for a gradient-based meta-learning mechanism. Furthermore, our analyses revealed a temporal hierarchy in learning: meta-level adaptation of the strategy operates on a slower timescale than learning action values.

Our discoveries were enabled by our novel `DynamicRL` method, designed to robustly track cognitive parameters over time, even when data are sparse and noisy, and it overcomes the static assumptions of traditional cognitive models. As a result, `DynamicRL` captures behavioral variability arising from time-varying strategies and strikes a balance between traditional interpretable symbolic models and “black-box” RNNs. It retains the normative and well-established structure of an RL model, while flexibly capturing dynamic strategy adaptation and avoiding the uninterpretable latent representations learned by RNNs directly fitted to behavior.

`DynamicRL` provides significant advantages in flexibility and generalizability over existing methods for modeling dynamic cognitive processes. Unlike naive sliding-window estimation (Sec. A.2), `DynamicRL` mitigates overfitting by estimating parameter dynamics jointly across participants and conditions, leveraging shared representations (Hinton et al., 2015). Task-specific hierarchical models rely on strong priors over generative processes (Mathys et al., 2011, 2014). Simulation-based (amortized) inference approaches (Schumacher et al., 2023; Ger et al., 2024), on the other hand, learn directly from synthetic data produced by predefined generative models, bypassing explicit likelihood evaluation. Thus, they do not directly quantify how well the model explains the observed data, and their validity depends critically on how accurately the predefined generative process reflects the true data-generating mechanism. In contrast, `DynamicRL` is trained directly on human behavioral data, imposing minimal assumptions about the generative process: humans use RL to update action values and make decisions.

Importantly, this methodological advance enables the exploration of a broader hypothesis space and opens new avenues for research, from modeling cognitive strategies to more nuanced temporal dynamics of strategy adaptation, thereby enabling powerful new applications in neuroscience and computational psychiatry. In cognitive neuroscience, these trial-level RL parameter estimates can serve as regressors in neural data analyses to identify meta-strategy encoding in neural activity or plasticity. Such findings would provide a critical link between behavioral-level computational models and their potential neural implementations.. In computational psychiatry, the framework provides a quantitative tool to investigate how abnormal meta-learning dynamics — such as excessive or insufficient strategy plasticity — may underlie conditions like anxiety, depression, or paranoia, offering a new means to characterize individual differences.

Future work could also generalize our approach to other cognitive models, such as the regression model, the drift-diffusion model (Ratcliff et al., 2016), or the temporal context model (Howard and Kahana, 2002), to track cognitive parameters, elucidate strategy adaptation dynamics, and further test the *gradient-based meta-learning* hypothesis. Moreover, systematic comparisons between different gradient-based and alternative optimization mechanisms (e.g., random search or heuristic rules) could provide further insights into the algorithmic principles underlying biological meta-learning. Our work also inspires the design of new experiments. For example, one could construct tasks that incorporate saddle points or plateaus in the reward landscape. Researchers can also study whether participants are capable of de novo exploitation of latent task structures to optimize the speed–accuracy trade-off (Xiong et al., 2025b).

Our work offers complementary behavioral support for the meta-RL framework in neuroscience (Wang et al., 2018), which conceptualizes the prefrontal cortex as a recurrent network that implements a learning algorithm trained through dopamine-driven meta-learning. Their framework presents a compelling hypothesis for how meta-learning may be implemented in the brain. Our approach begins with behavioral data and does not assume a predefined policy-gradient meta-learning structure, yet it reveals patterns of human strategy adaptation that resemble policy-gradient-driven meta-learning. Our work also resonates with longstanding multi-timescale learning theories in cognitive neuroscience (e.g., the complementary learning systems McClelland et al., 1995). Our results provide evidence for such dual-timescale learning, showing a fast process for learning action values alongside a slower meta-learning process that adjusts the learning strategy.

Our findings speak to a broader question: what computational principles unify biological and artificial adaptation? The gradient-based meta learning we observed parallel recent findings in large language models, which exhibit rapid “in-context learning” during inference. Recent work also shows that hierarchical meta-learning structures emerge within large language models to refine strategies over time (Coda-Forno et al., 2023). Together, these converging findings suggest that gradient-based meta-learning reflects a general principle of adaptive intelligence in biological and artificial systems.

## 4 Methods

### 4.1 Datasets

We analyze four human behavioral datasets collected from different bandit tasks with varying reward structures, including differences in environmental volatility, stochasticity, and horizon length. These tasks fall into two categories: restless bandit and stationary bandit. To facilitate model fitting and reward landscape simulations, rewards were normalized using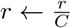, with task-specific constants *C* chosen to ensure comparable reward ranges where applicable.

#### Bahrami2020 (Four-armed restless bandit task)

In this task (Bahrami and Navajas, 2020), participants choose among four options whose mean rewards evolve stochastically over time according to an Ornstein–Uhlenbeck process centered at *µ* = 50, with a decay rate of *γ* = 0.9836 and process noise *σ*_OU_ = 2.8. At each time step, rewards are sampled as *r* ∼ 𝒩 (*µ, σ*^2^) with *σ* = 4. Each participant is required to complete *B* = 1 block of *T* = 150 trials. Due to timeouts or delayed responses, some participants completed fewer trials. 14 participants are removed from our analysis due to low performance (averaged raw reward lower than 43 points, which approximately corresponds to 1.5 standard deviations below the mean) and too many missing trials (fewer than 80 finished trials), resulting in *T*_total_ = 139,816 trials from *N* = 965 participants. Rewards were normalized using 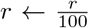, yielding a baseline mean of 0.5. To ensure data consistency for comparison, our analysis focused on the first 140 trials, which the majority of participants completed.

#### Suthaharan2021 (Three-armed probabilistic reversal learning task)

This task (Suthaharan et al., 2021) consists of *B* = 4 blocks of *T* = 40 trials each. Participants choose among three options with probabilistic rewards. In the first two blocks, reward probabilities are fixed at [0.9, 0.5, 0.1]; in the final two blocks, they change to [0.8, 0.5, 0.2]. After ten consecutive selections of the highest-reward option, reward contingencies reverse, enabling the study of behavioral flexibility in volatile environments. The dataset includes *N* = 1,012 participants and *T*_total_ = 173,120 trials. Because the baseline average reward is already 0.5, no normalization was applied.

#### Gershman2018 (Two-armed fixed-horizon bandit task)

This dataset is based on condition 2 of Gershman (2018), which features a symmetric task design. Condition 1, which is asymmetric because one option maintains a constant value throughout the experiment, was excluded for modeling simplicity. In each block, the expected rewards for the two arms are drawn from *µ*_*b*_ ∼ 𝒩 (*µ*, 10^2^), and actual rewards are sampled as *r* ∼ 𝒩 (*µ*_*b*_, 10). Participants complete *B* = 20 blocks of *T* = 10 trials each. The dataset includes *N* = 44 participants and *T*_total_ = 8,800 trials. Rewards were normalized via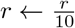, setting the baseline mean to 0.

#### Wilson2014 (Two-armed horizon task)

This task investigates exploration under uncertainty through a forced-choice phase followed by a free-choice phase (Wilson et al., 2014). In each block, participants choose between two arms: one arm’s mean reward is drawn from {40, 60}, and the other arm’s mean differs by an amount sampled from {4, 8, 12, 20, 30}. Rewards are sampled from a Gaussian distribution *r* ∼ 𝒩 (*µ, σ*^2^) with *σ* = 8. During the forced-choice phase (first 4 trials), actions are pre-assigned to create either symmetric (2 samples per arm) or asymmetric (3 vs. 1) information conditions. The subsequent free-choice phase varies in length, with horizons of either 1 or 6 trials. We aggregate data from *N* = 643 participants across six experiments (Wilson et al., 2014; Feng et al., 2021; Somerville et al., 2017; Waltz et al., 2020; Sadeghiyeh et al., 2020; Zajkowski et al., 2017). Of these, 34 participants were excluded if they met at least three of these criteria: low performance (average score below 52 raw points), highly asymmetric free-choice behavior (choosing the same option in more than 85% or exhibiting a choice autocorrelation greater than 0.85 or less than 0.15), and excessive fast responses (more than 20% of trials with reaction times under 30 ms). Due to differences in experimental design, the number of blocks (*B*) completed varies across participants, ranging from 300 to 1,200 free-choice trials, yielding a total of *T*_total_ = 754,890 trials. Rewards were normalized using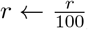, resulting in a baseline mean of 0.5. Although participants completed different numbers of blocks and encountered horizon conditions (1 or 6) in varying sequences, our DynamicRL model receives only subject identity, block number, and horizon condition as input. We restrict our analysis to blocks 0 through 60 for both horizon conditions to ensure comparability across participants. These early blocks are representative, as they were completed by the majority of participants.

### 4.2 DynamicRL method

#### 4.2.1 Model structures

In restless tasks, such as `Bahrami2020` and `Suthaharan2021`, the expected value of each option varies across trials within a block, requiring continuous adaptation. In this setting, we consider trial-level parameters *α*_*t*_ and *β*_*t*_ that define the RL strategy, allowing us to track strategy evolution over trials.

We use a gated recurrent unit (GRU) to estimate these RL parameters on a trial-by-trial basis. The model takes as input the previous action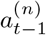, previous reward 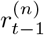, a learned participant embedding *E*^(*n*)^ (capturing individual differences), and trial-level contextual information 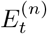. For `Bahrami2020`, 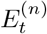 consists of the current trial number. For `Suthaharan2021`, 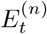 includes the current trial number, block number, and reward probability condition. The GRU updates its hidden state *h*_*t*_, which is passed to a linear readout *g*(·) to generate the RL parameters 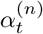 and 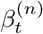 for participant *n* at trial *t*:

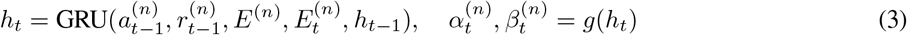

Unlike restless tasks, stationary tasks (`Gershman2018` and `Wilson2014`) assume that expected rewards of options are fixed within each block while varying across blocks. To focus on the slower inter-block timescale of meta-learning, we characterize the RL strategy across blocks with a single set of block-level RL parameters, *α*_*b*_ and *β*_*b*_. Under this formulation, model inputs are limited to block-level information (e.g., block number, horizon condition), excluding faster timescale, trial-level information (e.g., actions and rewards). By ignoring within-block variability, this formulation provides a parsimonious account of how a participant’s overall strategy adapts from one self-contained learning episode (a block) to the next.

We use a two-layer feedforward network (multilayer perceptron; MLP) to estimate RL parameters at the block level. The model receives the participant embedding *E*^(*n*)^ and block-level contextual information 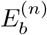. For `Gershman2018`, 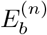 contains only the current block number. For `Wilson2014`, it includes the block number and the horizon condition. The MLP outputs the RL parameters 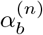and 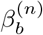 for participant *n* at block *b*:

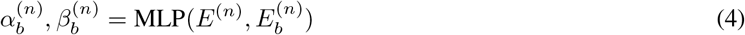

#### 4.2.2 Model fitting

To train our `DynamicRL` model, we use the negative log-likelihood (NLL) loss to maximize the likelihood of observed actions under the model’s predicted action probabilities. For a dataset with *N* participants and *T* trials per participant, the mean NLL loss is defined as:

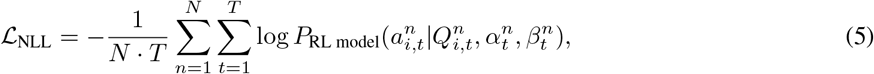

where 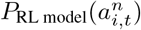 is the probability of choosing action *i* at trial *t* for participant *n*, as defined in Equation (2). The model parameters 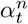 and 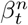 are estimated for each trial *t* in restless bandit tasks, or estimated for each block *b* in stationary bandit tasks.

To mitigate overfitting and ensure fair comparison across optimization methods, we employed a nested cross-validation protocol (Ji-An et al., 2025). Specifically, 70% of the data was used for training, 12.5% for validation (for early stopping and hyperparameter tuning), and 17.5% for testing. The outer loop used eight folds, and the inner loop used five.

Due to differences in task design across datasets, we tailored the cross-validation strategy accordingly. For `Gershman2018`, data were partitioned by block. For `Wilson2014`, data were segmented into non-overlapping 40-trial chunks. To preserve structural consistency, padding was introduced when needed, and padded trials were excluded from loss computation. For `Bahrami2020` and `Suthaharan2021`, trials were split into non-overlapping sets using distinct masks for training, validation, and testing (i.e., an interspersed split protocol Ji-An et al., 2025; Hans et al., 2024).

To improve robustness during training, we added Gaussian noise 𝒩 (0, 0.01) to reward values. Early stopping was applied if validation performance did not improve for 20 consecutive epochs. We used the AdamW optimizer with default settings, a batch size of 64, and a participant embedding size of 20. Other hyperparameters were selected via grid search over the following values: learning rate in {0.005, 0.01, 0.02, 0.03}, weight decay in {0.01, 0.1}, dropout in {0, 0.1, 0.2}, and hidden layer size *h* ∈ {50, 100}. To constrain RL parameter estimates, we applied a sigmoid activation to ensure *α* ∈ [0, 1], and a ReLU activation to enforce *β* ≥ 0. The best model was selected based on the lowest average NLL on the test data across all folds.

### 4.3 Comparing early and late task performance

To assess whether participants improved their decision-making performance over time, we examined a metric termed *advantage*, defined as the difference between the reward obtained by a participant and their average reward. To ensure comparability across datasets with differing reward structures, advantage scores were normalized: for `Gershman2018`, the baseline was set to 0; for the remaining datasets, it was set to 0.5. This normalization was intended to reduce cross-subject variance arising from heterogeneous reward trajectories.

For datasets without distinct experimental conditions, we conducted paired *t*-tests to compare early versus late task performance. For datasets with within-subject manipulations, we employed linear mixed-effects models to account for repeated measures. Specifically, for `Suthaharan2021`, we tested whether performance in the final 20 trials exceeded that in the first 20 trials using the following model:

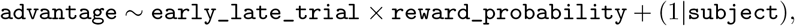

implemented using statsmodels.MixedLM (Seabold and Perktold, 2010). For `Wilson2014`, we examined whether participants performed better in later blocks with:

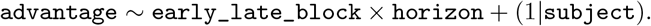

In `Bahrami2020` and `Wilson2014`, not all participants completed enough trials or blocks. To ensure the robustness and interpretability of the model estimates, we included only task segments with sufficient data coverage across participants.

### 4.4 Reward landscape

To understand the task performance of different RL parameters combinations under each task, we constructed reward landscapes by simulating an RL agent with varying combinations of RL parameters. For each task, we systematically varied the learning rate *α* from 0 to 1 in increments of 0.025. The inverse temperature parameter *β* was varied across a broader range (0 to 60) using non-linear increments in three regions: 0 to 16 in steps of 0.5, 16 to 32 in steps of 1, and 32 to 60 in steps of 3, enabling finer resolution in regions of high sensitivity to *β*. This process yields a total of 1,302 parameter combinations. For each combination, we simulated behavior over 200 blocks of 120 trials and recorded the resulting average reward *R*. We then plotted reward landscapes *R* as a function of (*α, β*).

To ensure smooth visualization, we applied cubic interpolation to produce a dense grid with 100 points for *α* (from 0 to 1) and 200 points for *β* (from 0 to 60). To reduce high-frequency noise that arises from finite simulations, a two-dimensional Gaussian filter with a standard deviation of 3 was applied to the interpolated reward landscapes.

For tasks with multiple experimental conditions, we display representative conditions. Specifically, for `Suthaharan2021`, we show the reward landscape for the first two blocks under the [0.9, 0.5, 0.1] reward probability condition; the landscape under the [0.8, 0.5, 0.2] condition exhibits similar characteristics. For `Wilson2014`, we present results for the horizon-6 condition, as the horizon-1 condition produces comparable reward landscapes.

### 4.5 Alignment of human RL parameter updates with the reward gradient

The alignment of human RL parameter updates with the reward gradient was calculated at the trial level for `Bahrami2020` and `Suthaharan2021` and at the block level for `Gershman2018` and `Wilson2014`.

#### 4.5.1 Cosine similarity with the reward gradient in parameter space

In artificial agents, policy gradient ascent typically requires averaging over many repeated episodes to reduce the variance of policy gradient estimation. However, each human participant completed only a single action-outcome sequence per task. Because this limited data naturally results in high variance in estimated gradient signals, we calculate the ‘optimal gradient vector’ from the simulated reward landscape rather than from a noisy empirical gradient to ensure a stable comparison across participants.

We first computed the cosine similarity between participants’ RL parameter updates and the corresponding gradient direction on the reward landscape *R*(*α, β*). Specifically, the gradient of the reward surface was calculated using finite differences:

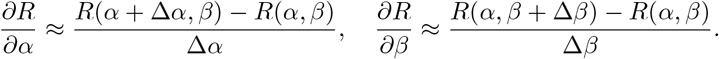

This yields a gradient vector field 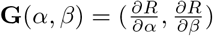 that estimates the direction in the parameter space along which the expected obtained reward is maximally increased.

For each participant *s*, we denote their RL parameters at time *t* (either a trial or a block) as 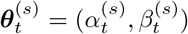. The parameter update vector is:

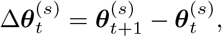

and we evaluate the reward gradient 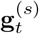 at the midpoint of each segment:

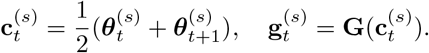

We then compute their cosine similarity:

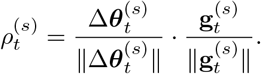

The overall alignment score for participant *s* is defined as the mean cosine similarity across all valid segments:

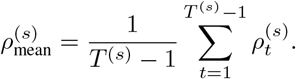

This metric quantifies the degree to which human RL parameter updates follow the optimal gradient direction. A value near 1 indicates strong alignment, while values near 0 or negative suggest deviation from the reward-maximizing direction. Perfect alignment is not expected due to the stochasticity inherent in online learning, but positive alignment at the group level provides evidence for gradient-like updates.

#### 4.5.2 Optimality of meta-learning update in reward space

To provide a measure of alignment that is invariant to the scale of parameters, we also evaluated whether participants’ RL parameter updates shift parameters toward regions with higher expected reward in the parameter space.

Using the precomputed reward landscape *R*(*α, β*), we estimated the expected rewards associated with each participant’s RL parameters at different time points. For participant *s* at time *t* (trial or block), the estimated parameters 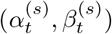 were mapped to their corresponding expected rewards using a nearest-neighbor lookup on a 100 *×* 200 grid:

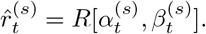

This yielded a reward trajectory 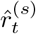 over time for each participant.

To assess whether these trajectories reflect systematic improvement, we conducted a lagged effect analysis. For each participant and a time lag Δ*t* (either at the trial or block level), we computed Cohen’s *d* (i.e., differences normalized by standard deviations) between expected rewards 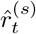 at time *t* and 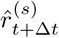 :

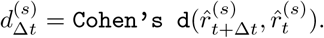

A positive *d*_Δ*t*_ indicates that RL parameter updates tend to increase expected reward, consistent with a reward-directed optimization process.

Together, these complementary measures—cosine similarity in the parameter space and reward improvement in the outcome space—allow us to rigorously assess whether human strategy updates are aligned with gradient directions.

### 4.6 Correlation between gradient SNR and RL parameter strategy improvement

Here, we examined whether the reliability of the reward gradient along the estimated RL trajectories predicts the magnitude of RL parameter updates. Specifically, we asked whether regions of the reward landscape with more reliable gradient signals—quantified as a noise-normalized gradient magnitude, or gradient SNR—are associated with larger step sizes in participants’ parameter trajectories.

#### 4.6.1 Computation of the noise-normalized gradient 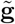

For each participant *s* and time point *t*, we first computed the finite-difference approximation of the reward gradient:

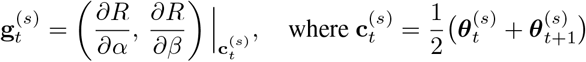

denotes the midpoint between consecutive parameter states 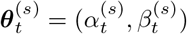 and 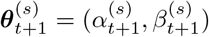. This pairing procedure ensures that each gradient estimate 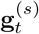 corresponds directly to a single parameter update between consecutive trials or blocks.

The gradient components were computed using finite differences on the reward landscape *R*(*α, β*) estimated for each participant. For interior grid points, we applied central differences:

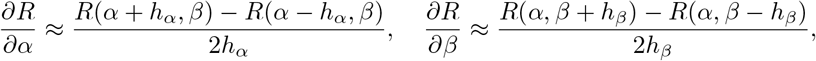

and used one-sided differences for boundary points.

To assess the reliability of these gradient estimates, we propagated the empirical variance of the reward estimates through the finite-difference stencil. Specifically,

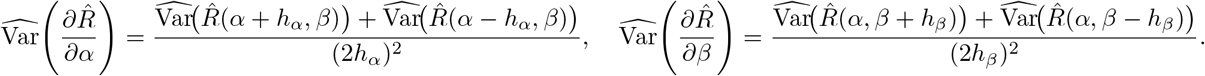

Here, 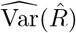 denotes the empirical variance of the estimated expected reward at each grid point, computed from 100 independent simulated trajectories. Since each simulation used independent random seeds, covariance terms between neighboring grid points were omitted.

Under these assumptions, the noise-normalized gradient magnitude (gradient SNR) was defined as:

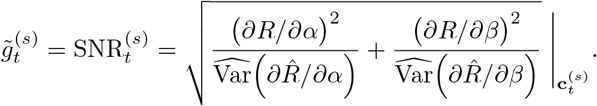

Higher values of 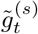 indicate stronger and more reliable gradients, corresponding to regions of the reward landscape where optimization signals are both large and consistent across stochastic realizations.

### 4.6.2 Correlation with step-size

For each participant *s* and time point *t*, we quantified the magnitude of the corresponding parameter update as the step size in parameter space:

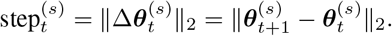

Each gradient–update pair 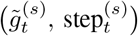 thus reflects how strongly and reliably the local reward gradient predicted the magnitude of the participant’s adjustment in parameter space.

We then evaluated the relationship between gradient reliability and update magnitude by computing the Pearson correlation between 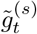 and 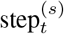 across all participants and time points. A positive correlation would indicate that more reliable gradients tend to drive larger parameter adjustments, suggesting sensitivity of the learning dynamics to the reliability of the underlying optimization signal.

#### 4.6.3 Statistical significance assessment

To assess statistical significance, we performed permutation tests with 10,000 iterations. Within each participant, we randomly permuted the temporal correspondence between 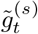 and the target variable along the trial or block dimension to construct a null distribution of correlation coefficients. The empirical correlation was compared to this null distribution to obtain p-values. All computations of 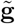 were based on the same dense, smoothed reward landscapes described in Section 4.4.

### 4.7 Autocorrelation and timescale

We computed autocorrelation for each participant using the standard implementation from `statsmodels.tsa.stattools.acf`. Due to differences in trial count per block, we set the maximum number of time lags *T* to 20 for `Bahrami2020` and 4 for `Suthaharan2021`. We then averaged the autocorrelation across participants to produce group-level autocorrelation *ρ*(*t*) at lag *t*.

To quantify the effective timescale of each variable, we computed the integrated autocorrelation time (Sokal, 1997):

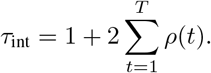

Larger *τ*_int_ indicates greater temporal persistence and slower variation in the corresponding variable.

## 5 Acknowledgments

We thank the researchers who generously shared their datasets. This work was primarily conducted while HDX and RCW were at the University of Arizona and was supported by NIH grant R01AG061888 awarded to RCW. We also acknowledge the University of Arizona’s High Performance Computing (HPC) facilities for providing essential computational resources.

## 7 Data Availability

The horizon task, `Wilson2014` dataset (Wilson et al., 2014), is available upon request. The two-armed bandit task, `Gershman2018` dataset (Gershman, 2018), was sourced from task2 in and is available at https://github.com/sjgershm/exploration. The three-armed probabilistic reversal learning task, `Suthaharan2021` dataset (Suthaharan et al., 2021), was sourced from and is available at https://github.com/psuthaharan/covid19paranoia. The four-arm restless bandit task, `Bahrami2020` dataset (Bahrami and Navajas, 2020), is available at https://osf.io/f3t2a/.

## 8 Author Contributions

H.D.X., L.J.A. conceived the study and drafted the initial manuscript. H.D.X. created and applied the models, conducted the data analysis. R.C.W. provided funding support. R.C.W., M.G.M supervised the project. All authors contributed to revising the manuscript and approved the final version.

## A Supplementary Material

### A.1 Dynamic RL parameter recovery

To evaluate the validity and robustness of the `DynamicRL` framework, we conducted a model recovery analysis using synthetic behavioral data generated from RL agents with predefined dynamic RL parameter trajectories. This analysis assesses whether the model can reliably recover the temporal dynamics of RL parameters when the ground-truth values are known (Wilson and Collins, 2019).

We tested model recovery on two datasets with trial-level dynamics: `Bahrami2020` and `Suthaharan2021`. In `Bahrami2020`, we used the original 4-armed restless bandit task without controlling for the three identical reward schedules. For `Suthaharan2021`, we focused on the condition with reward probabilities [0.9, 0.5, 0.1].

For each dataset, we defined a dynamic range of *α* and *β*, and generated linear RL parameter trajectories (increasing, decreasing, or constant) across trials. Initial and final parameter values were sampled from a 3-by-3 grid within empirically observed ranges: [0.4, 0.8] for *α* in both datasets, [4, 10] for *β* in `Bahrami2020`, and [2, 8] for *β* in `Suthaharan2021`. Each condition was repeated ten times using different synthetic agent IDs, simulating distinct participants employing the same strategies and producing a diverse set of RL parameter trajectories. The parameter ranges were selected based on the interquartile range (second to third quartile) of estimates from the fixed-parameter models and were slightly expanded to ensure broader coverage.

We then applied the `DynamicRL` model to these synthetic datasets, using the same fitting and selection procedure described in Sec. 4.2.2, and compared the recovered trajectories to the ground-truth values. In both datasets, the model successfully captured the qualitative trend of parameter evolution. Constant parameters were recovered as stable, flat trajectories. Monotonic changes in *α* and *β* were consistently reflected in the recovered trajectories (Fig. A.1, A.2). Quantitatively, recovered trajectories were strongly correlated with the true values. Across conditions, most Pearson correlations exceeded 0.95. Recovery accuracy was slightly lower in `Suthaharan2021`, likely due to fewer trials and discontinuities introduced by block structures. Nonetheless, overall parameter recovery performance remained high, demonstrating that DynamicRL effectively identifies dynamic RL parameters under a broad range of conditions.

### A.2 Alternative moving average RL model

A natural baseline for estimating time-varying RL parameters is the sliding window approach: partition the trial sequence into overlapping windows and fit standard RL models within each window. This method is conceptually simple and can, in principle, capture non-stationary changes in behavior over time.

To empirically evaluate this approach, we applied a sliding-window RL estimator to the `Bahrami2020` dataset, which contains over 100 trials per participant. We selected this dataset because the relatively long trial sequences permit the use of larger window sizes. Other datasets were excluded due to insufficient trial counts (`Suthaharan2021` has 40 trials) or block counts (`Gershman2018` has 20 blocks; `Wilson2014` has 60 blocks), which would restrict window size and reduce the reliability of parameter estimation—particularly given the stochastic nature of rewards and the heightened risk of overfitting when data within each window are sparse.

We systematically varied the window size, using a window stride of 5 trials (e.g., for a window size of 30, windows covered trials 0–30, 5–35, etc.). Within each window, we fit fixed RL parameters using maximum likelihood estimation with the Nelder–Mead optimization method, initialized from 16 grid-based starting points. We constructed this grid by uniformly sampling four values for each parameter: *α* ∈ [0, 1], *β* ∈ [0, 60]. This initialization strategy established a strong baseline for comparison.

Because a principled train-test split is not straightforward in this setting, we directly compared the training negative log-likelihood (NLL) of the sliding-window model to the test NLL of both the standard fixed-RL model and the `DynamicRL` model. We notice that the training NLL usually exaggerates the predictive performance due to overfitting.

As shown in Fig. A.3, small windows (e.g., 10 trials) yield lower training NLL but produce noisy estimates and are potentially prone to overfit since it leads to higher variance in parameter estimation. Further, these still perform worse than the test loss of `DynamicRL`. Larger windows improve stability but fail to capture fine-grained trial-level dynamics, sacrificing temporal resolution. Even at optimal window size, the sliding-window model underperforms relative to `DynamicRL`.

Therefore, the neural network architecture in `DynamicRL` is necessary to provide robust and accurate estimation by learning to infer RL parameters across the full dataset. It leverages shared strategies across participants and conditions, producing smooth and context-sensitive parameter trajectories that generalize well.

**Figure A1.**
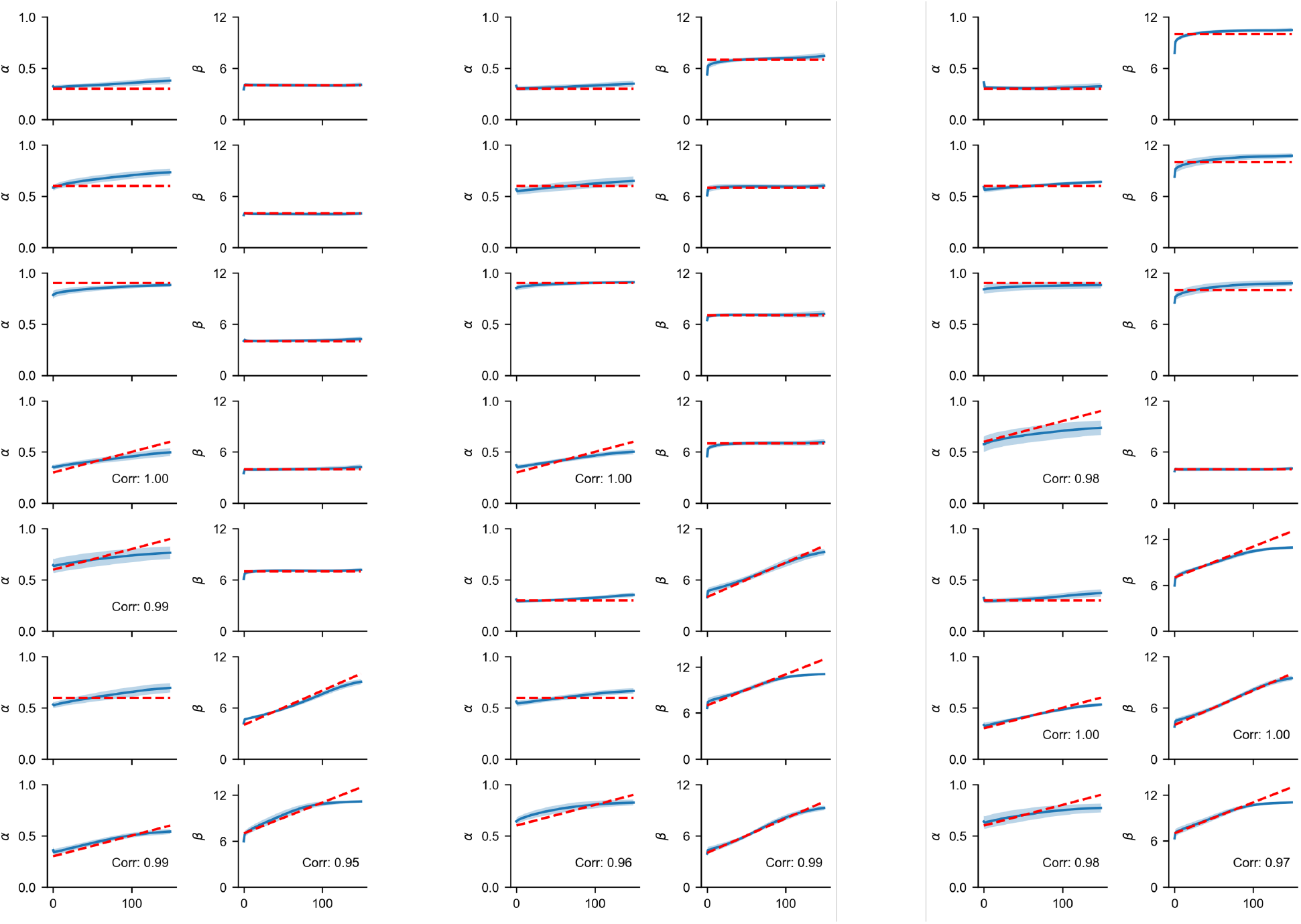
Dynamic RL parameter for Bahrami2020. Red dashed lines indicate ground-truth RL parameter values; blue lines are estimates by DynamicRL from data.

### A.3 Supplementary figures

**Figure A2.**
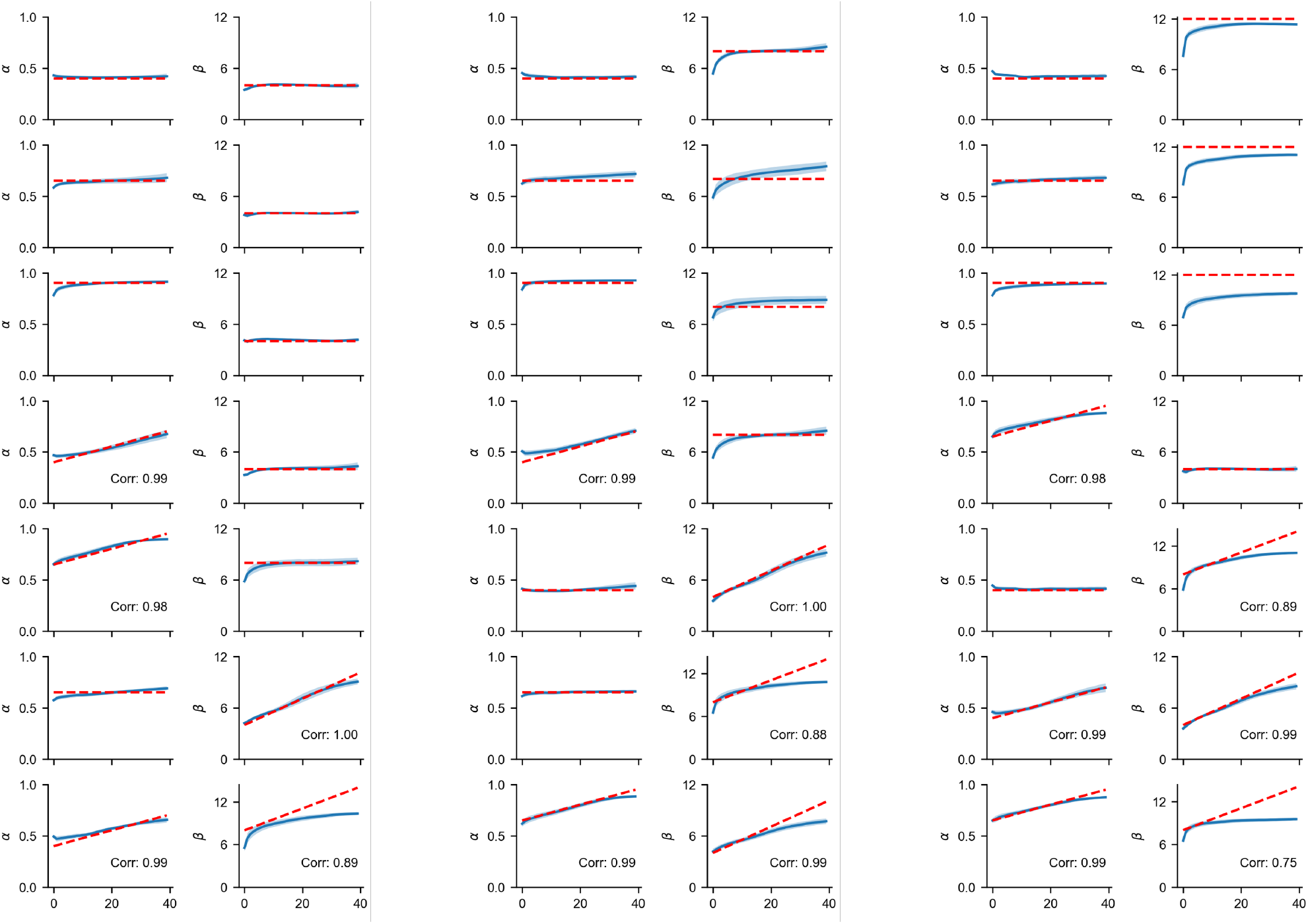
Dynamic RL parameter recovery for Suthaharan2021. Red dashed lines indicate ground-truth RL parameter values; blue lines are estimates by DynamicRL from data.

**Figure A3.**
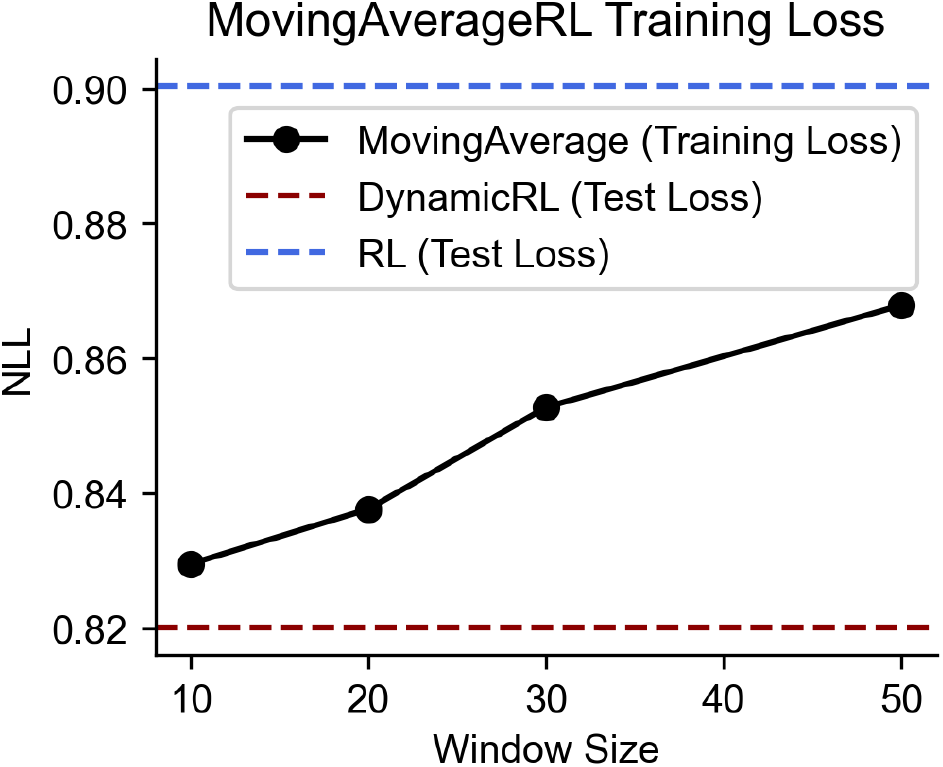
Performance of sliding-window RL estimation on Bahrami2020. Training NLL increases with window size (i.e., fit performance decreases). Smaller windows overfit and produce unstable estimates. Even the best-performing sliding-window model underperforms relative to DynamicRL on held-out data.

**Figure A4.**
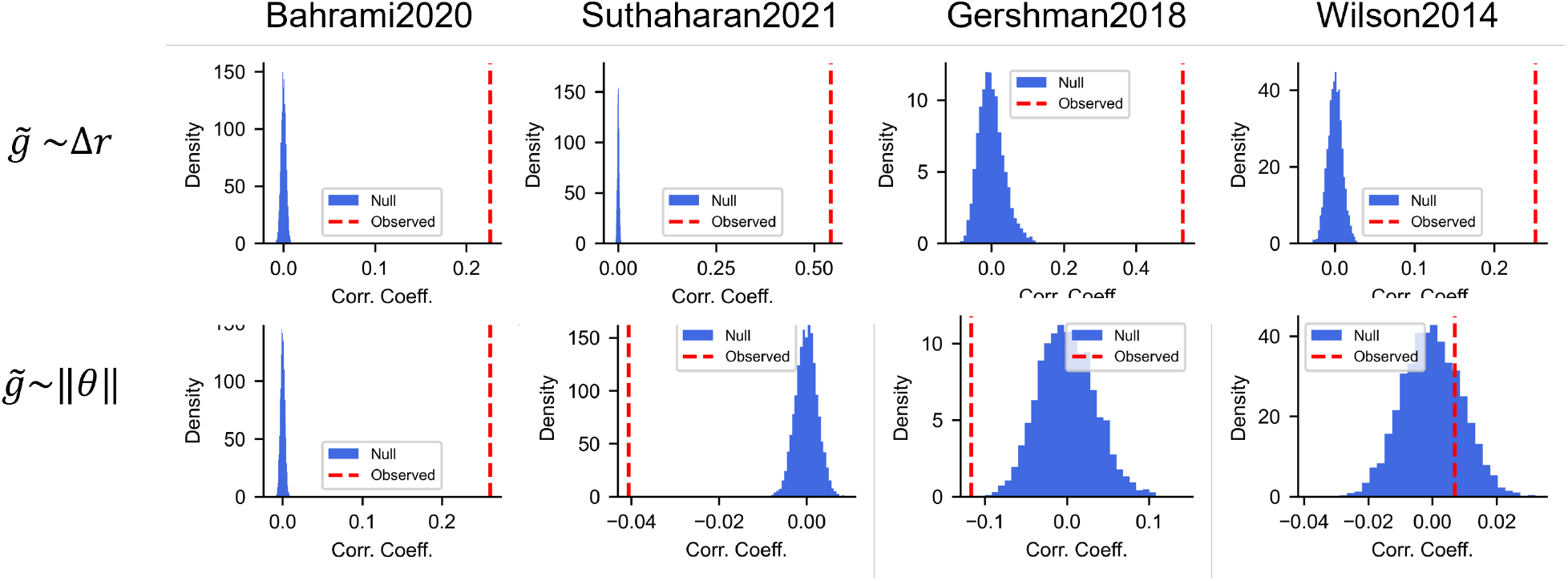
Correlation between gradient norm and meta-learning strategy improvement. For each dataset, we computed the empirical correlation between the magnitude of the local reward gradient and the optimality of meta-learning updates. The blue histogram shows the null distribution obtained via permutation testing. The red dashed line indicates the observed correlation coefficient. In all datasets, the observed correlation significantly exceeds the null distribution.

